# RiceLncPedia: a comprehensive database of rice long non-coding RNAs

**DOI:** 10.1101/2020.05.22.110569

**Authors:** Zhengfeng Zhang, Yao Xu, Fei Yang, Benze Xiao, Guoliang Li

## Abstract

Long non-coding RNAs (lncRNAs) play significant functions in various biological processes including differentiation, development and adaptation to different environments. Although multi research focused on lncRNAs in rice, the systematic identification and annotation of lncRNAs expressed in different tissues, developmental stages under diverse conditions are still scarce. This impacts the elucidation of their functional significance and the further research on them. Here, RiceLncPedia (http://218.199.68.191:10092/) is constructed including rice lncRNAs explored from 2313 publically available rice RNA-seq libraries and characterize them with multi-omics data sets. In the current version, RiceLncPedia shows 6978 lncRNAs with abundant features: (i) expression profile across 2313 rice RNA-seq libraries; (ii) an online genome browser for rice lncRNAs; (iii) genome SNPs in lncRNA transcripts; (iv) lncRNA associations with phenotype; (v) overlap of lncRNAs with transposons; and (vi) LncRNA-miRNA interactions and lncRNAs as the precursors of miRNAs. In total, RiceLncPedia imported numerous of rice lncRNAs during development under various environments as well as their features extracted from multi-omics data and thus serve as a fruitful resource for rice-related research communities. RiceLncPedia will be further updated with experimental validation, functions association and epigenetic characteristics to greatly facilitate future investigation on rice lncRNAs.

## INTRODUCTION

Long noncoding RNAs (lncRNAs) are referred as RNA molecules with length of at least 200 nucleotides (nt) and usually have low protein coding potential (1). In human, function of lncRNAs is relevant to various important biological processes such as cell differentiation, immune response, diverse cancers and so on (2-7). In plants, emerging evidences indicate that lncRNAs function as key modulators in a wide range of biological processes including development and stress response at the epigenetic, transcriptional and post-transcriptional levels (8-16).

In recent years, the explosive high-throughput sequencing promotes the global discovery of lncRNAs in various processes in animal system as well as plants. Accordingly, multiple databases have been constructed, focusing on different aspect of lncRNAs in human and various animal species. NONCODE 5.0 provides the gene expression pattern from RNA-seq data and includes lncRNAs in 17 species such as human, mouse and so on (17). LncRNAdb v2.0 (18) and LNCipedia (19) are annotation databases based on literature evidence. LncRNAs with the support of coexpression, differential expression, binding proteins and phylogenetic conservation was constructed in database of lncRNAtor (20). lncRNASNP integrated the lncRNA, SNP, GWAS results and miRNA expression profiles in human and mouse (21). Recently, a comprehensive database of human long non-coding RNAs, LncBook was constructed which incorporated multi-omics data including expression profile, sequence variation, association with miRNA, epigenetic features and diseases association to annotate human lncRNAs (22). Several lncRNAs databases were also developed in plants but mainly for model species *Arabidopsis* (23,24). Compared with human and animal lncRNA databases, the number of plant lncRNA databases and lncRNA features referred in the databases were relatively rare. Those plant databases continuously accessible and updated including rice lncRNAs were even fewer (Table 1 Key differences between the existing databases including rice lncRNAs and RiceLncPedia). A long non-coding RNA database of plants (PLncDB) provides rich features for lncRNAs including genomic information, expression profiles in multi developmental stages, mutants and stress conditions, epigenetic modification and small RNA associations (25). However, this database only focused on Arabidopsis without other plant species in the present version. GREENC, a Wiki-based database of plant lncRNAs, identified 120 000 lncRNAs from 37 plant species and six algae, including 5237 lncRNAs from rice (26). The GreeNC database provides information about sequence, genomic coordinates, coding potential and folding energy for all the identified lncRNAs. However, the lncRNAs in this database were identified from the transcripts in Phytozome v10.3 and therefore the expression profile across tissues, growth conditions were unavailable. Additionally, variations and other genomic features are lack either (26,27). CANTATAdb collected lncRNAs in 36 plant species and 3 algae, among of which, 2788 lncRNAs are collected from only 8 rice RNA-Seq libraries (28). PNRD, a plant non-coding RNA database, collected a total of 25739 entries of 11 different types of ncRNAs from 150 plant species, whereas only harbors 790 lncRNAs in rice (29). PLNlncRbase, A resource for experimentally identified lncRNAs in plants, has manually collected data from nearly 200 published literature, covering 1187 plant lncRNAs in 43 plant species, 1060 of which are stress-related lncRNAs under 17 different abiotic or biotic stress conditions in various plant species (30). Another database including low-throughput experiment validated lncRNAs is EVLncRNAs, which contains 1543 lncRNAs from 77 species, whereas only 428 plant lncRNAs from 44 plant species (31). PLncPRO has discovered lncRNAs responsive to abiotic stresses in rice and chickpea, where nine rice RNA-seq samples were explored, associating with desiccation and salinity stresses from only three rice cultivars (32). However, these databases are small in scale and less comprehensive for rice, a most widely planted staple food crop and model crop. A comprehensive rice database covering lncRNAs from more widely developmental tissues, stages and diverse stress condition as well as integrating multi-omic features is still lacking (Table 1).

**Table 1.** Key differences between the existing rice lncRNA databases and RiceLncPedia.

| Data Item | RiceLncPedia | GREENC | CANTATAdb 2.0 | PNRD | PLNlncRbase | PLncPRO | EVLncRNAs |
| --- | --- | --- | --- | --- | --- | --- | --- |
| Species | **Rice** | 43 species | 39 species | 150 species | 43 species | Rice, chickpea | 77 |
| LncRNA entries | **6978** | 120,000 | 239, 631 | 25739 ncRNAs | 1187 | 12314 | 1543 |
| Rice LncRNA entries | **6978** | 5237 | 2788 | 790 | unknown* | 7345 | 40 |
| Rice RNA-seq libraries | **2313** | NA | 8 | NA | NA | 9 | NA |
| Experimentally identified lncRNAs |  | NA | NA | Partial | 1060 | NA | √ |
| Sequences | √ | √ | √ | √ | √ | √ | √ |
| coding potential | √ | √ | √ | √ | NA | NA | NA |
| Genomic information | √ | √ | √ | √ | √ | NA | √ |
| Expression | √ | NA | √ | NA | √ | NA | √ |
| Small RNA associations | √ | √ | NA | NA | NA | NA | NA |
| LncRNA genome variation | √ | NA | NA | NA | NA | NA | NA |
| Phenotype association | √ | NA | NA | NA | √ | NA | √ |
| TE-lncRNAs | √ | √ | NA | NA | NA | NA | NA |
| Accessible | √ | √ | √ | √ | NA | √ | √ |

Note: \* represents the unknown number of rice lncRNAs since the database is not accessible following the link in the reference.

In this study, we developed a rice lncRNAs database, RiceLncPedia with the following comprehensive features, attempting to facilitate the understanding and usage of rice lncRNAs: (i) a collection of lncRNAs identified from the majority of the publically available rice RNA-seq dataset (2313 RNA-seq libraries); (ii) lncRNA expression profiles in various tissues, developmental stages, stress treatments and different cultivars; (iii) lncRNA associations with genome variations; (iv) the linkage of lncRNAs with phenotype (inferred from public available QTLs and GWAS results); (v) the overlap information of lncRNAs and transposon elements, since the transposon elements (TEs) play key roles in the generation and function of lncRNAs; (vi) the lncRNAs predicted as miRNA targets or miRNA precursors collected as well.

## MATERIALS AND METHODS

### Data collection

We identified rice lncRNAs from RNA assemblies based on 2313 public available RNA-seq libraries. To obtain high-confidence lncRNAs, the following steps were adopted from the raw RNA-seq data(Figure 1). Low-quality reads with >5% ambiguous bases were filtered and adapter sequences were trimmed. The clean RNA-seq reads were mapped to the rice reference genome Os-Nipponbare-Reference-IRGSP-1.0 using Hisat 2.1.0 program with the default parameters set (33). Transcriptomes were reconstructed using StringTie v1.3.3 package with the default parameters (33). Transcriptome assemblies generated from the above steps were subsequently merged with StringTie --merge to acquire comprehensive non-redundant transcripts for subsequent analysis: (i) the transcripts assembled using StringTie were then annotated with Cuffcompare program to filter out known protein-coding transcripts, rRNA and tRNA with the comparison code ‘=’; questionable transcripts tagged with codes ‘e’, ‘p’ and ‘s’ by Cuffcompare were filtered out; (iii) transcripts with lengths more than 200nt were selected as lncRNA candidates;(iv) transcripts with FPKM scores smaller than 0.5 in all samples were discarded; (v) further screened through the protein-coding score test using Coding Potential Calculator (CPC2) (34), Plant Long Non-Coding RNA Prediction by Random fOrests (PlncPRO) (32) and Protein family database (Pfam) (35). As a consequence, a total of 6978 non-redundant rice lncRNA transcripts were obtained, belonging to 5845 gene loci.

**Figure 1.**
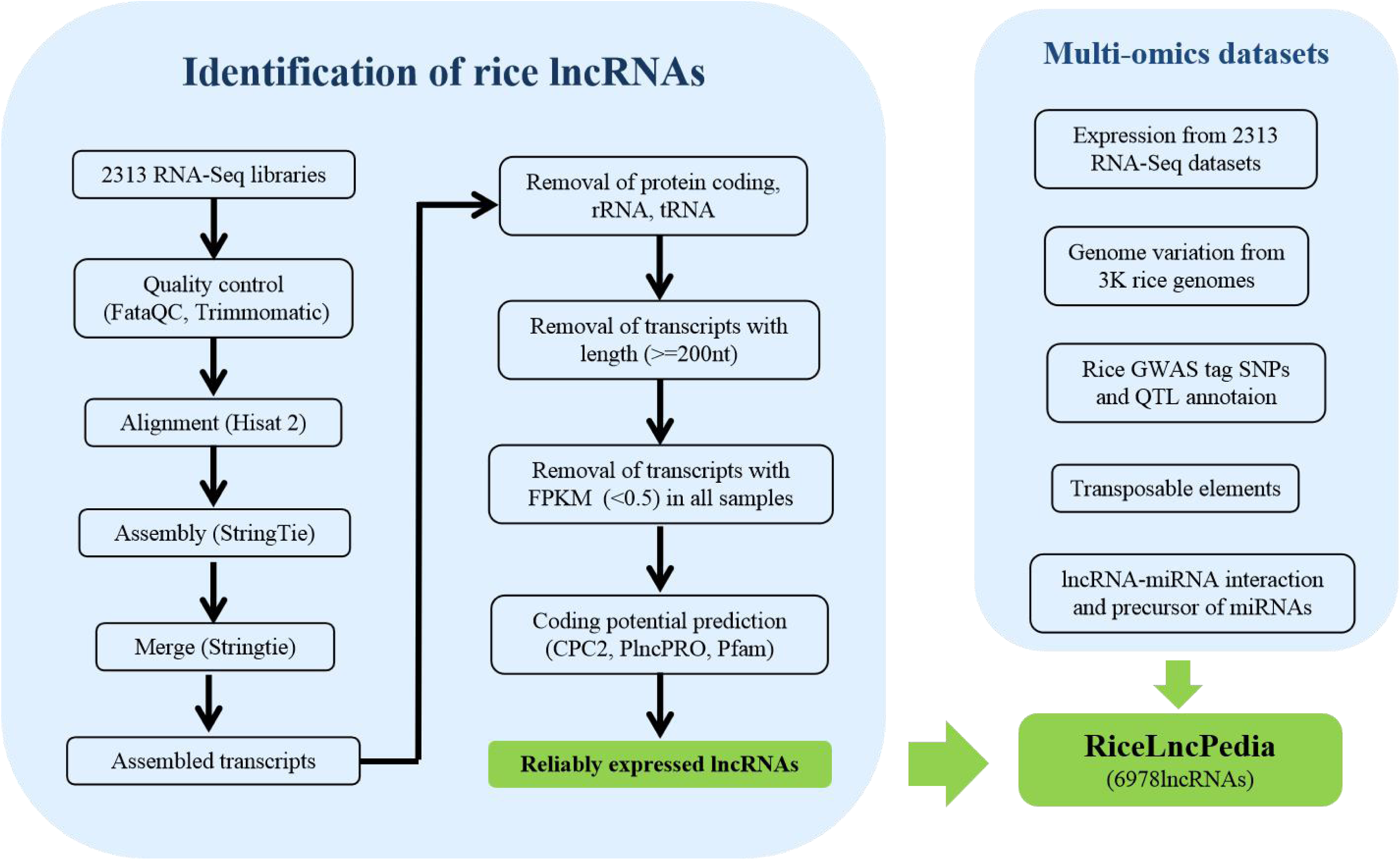
Pipeline for rice lncRNAs identification and multi-omics data integration.

### Data integration and annotation

#### 1) Molecular features of rice lncRNAs

Based on the location of lncRNAs relative to proteincoding RNAs, the different types of lncRNAs include intergenic lncRNA (lincRNA), intronic lncRNA, sense lncRNA, antisense lncRNA, which were tagged using Cuffcompare(36)as class code of u, i, o and x, respectively. As code ‘j’ represents potentially novel isoform: at least one splice junction is shared with a reference transcript (Cuffcompare manuscript), this class of transcripts can be long non-coding isoforms of known genes (37). Thereby, the transcripts with code of u, i, j, o or x were retained for further analysis. The number, length and GC content (%) of lncRNAs were counted by in-house Python scripts.

#### 2) LncRNAs expression

StringTie 1.3.3 was used to calculate FPKMs of lncRNAs. Housekeeping (HK) lncRNAs, tissue-specific (TS) and stress responsive (SR) lncRNAs were determined as following (22) with minor modification. Briefly, based on the expression value of lncRNAs in the selected datasets (see the section of DATABASE CONTENT AND FEATURES, Expression profile), τ-value and coefficient of variance (cv) were applied to distinguish HK lncRNAs (τ-value ≤ 0.5 and cv ≤ 0.5) and SR (stress response) or TS (tissue specific) lncRNAs (τ -value ≥ 0.95). Here the index τ was defined as: 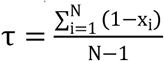, where N stands for the number of tissues and x_i_ represents the expression profile component normalized by the maximal component value (22, 38). To explore the relationship of sample grouping with lncRNA expression profiles, we selected 339 libraries (Table S1) with clearly tissue, variety and treatments information and drew the hierarchical clustering heatmap based on lncRNAs expression FPKMs using R package pheatmap1.0.12 (http://cran.r-project.org/web/packages/pheatmap/index.html).

#### 3) LncRNA genome variation

Single nucleotide polymorphism (SNP) genotyping data were downloaded from 3000 rice genome Projects (http://snpseek.irri.org/_download.zul) (39), which were called against the reference genome Os-Nipponbare-Reference-IRGSP-1.0. The SNP mapped to any lncRNA site (from the start to the end position of lncRNA on the genome) were retrieved as the lncRNA-SNP.

#### 4) Association of LncRNAswith agricultural traits

Two datasets were applied to predict the association of lncRNAs with agricultural traits. GWAS tag SNPs were downloaded from Rice SNP-Seek database (40) and QTLs information from Q-TARO (QTL Annotation Rice Online) database (41). The GWAS SNP contributed phenotype was predicted as possible function of a specific lncRNA if that GWAS tag SNPs co-located with this lncRNA. Accordingly, when a specific lncRNA locus overlapped with a QTL, this QTL-related trait was tagged on this lncRNA as a predicted phenotype association.

#### 5) LncRNA-miRNA interaction and precursor of miRNAs

psRNA Target was used to predict the lncRNA target of microRNAs with the default parameters. This procedure screened candidate interactions of lncRNA–miRNAs (42). The precursors of miRNAs were screened by comparing lncRNAs sequences with rice pre-miRNAs hairpin sequences (http://www.mirbase.org/). Blast 2.7 was used with the threshold e-value ≤ 10−5, coverage percent bigger than 90% and -max_hsps as 1.

#### 6) Transposon-lncRNA

Genomic coordinates of Japonicatransposon elements (TE) were downloaded from https://www.genome.arizona.edu/cgi-bin/rite/index.cgi(43). The position of TE was compared with respect to lncRNAs in rice genome. LncRNAs overlapping with TE were characterized as TE-lncRNAs associations.

### Implementation

We constructed RiceLncPedia database using Django as back end web framework and PostgreSQL(https://www.postgresql.org/) as database engine. JQuery and AJAX (Asynchronous JavaScript and XML) were used to developed web interfaces. As for front-end framework, we employed Bootstrap (https://getbootstrap.com) to supply a series of templates to design web pages with consistent interface components. Additionally, we adopted the icon in Font Awesome in RiceLncPedia website (http://www.fontawesome.com.cn/). Data visualization was powered by Pyecharts (https://github.com/pyecharts/pyecharts) to add interactive diagrams to our website.

## DATABASE CONTENT AND FEATURES

In contrast with the existing lncRNA database, the present database features a comprehensive collection of rice lncRNAs from most widely samples and systematic curation of lncRNA annotation through integrating multi-omics data, covering molecular features, expression profiles, sequences features and agricultural traits association.

### Number of lncRNA transcripts

RiceLncPedia accommodates 6978 rice lncRNAs which were identified based on 2313 RNA-seq libraries analysis, belonging to 5845 gene loci. The lncRNAs were organized in RiceLncPedia as transcripts. Each lncRNA transcript entity is assigned to a unique accession number and a specific page was linked to each lncRNA transcript which shows detailed molecular features (Transcript id; Location; Classification; Length; GC Content (%); Exon Number; Sequence; Coding Potential; Genome Browser), Genome Variation, Transposon elements, Small RNA targets, miRNA precursors and expression profile across represented RNA-seq samples (Figure 2).

**Figure 2.**
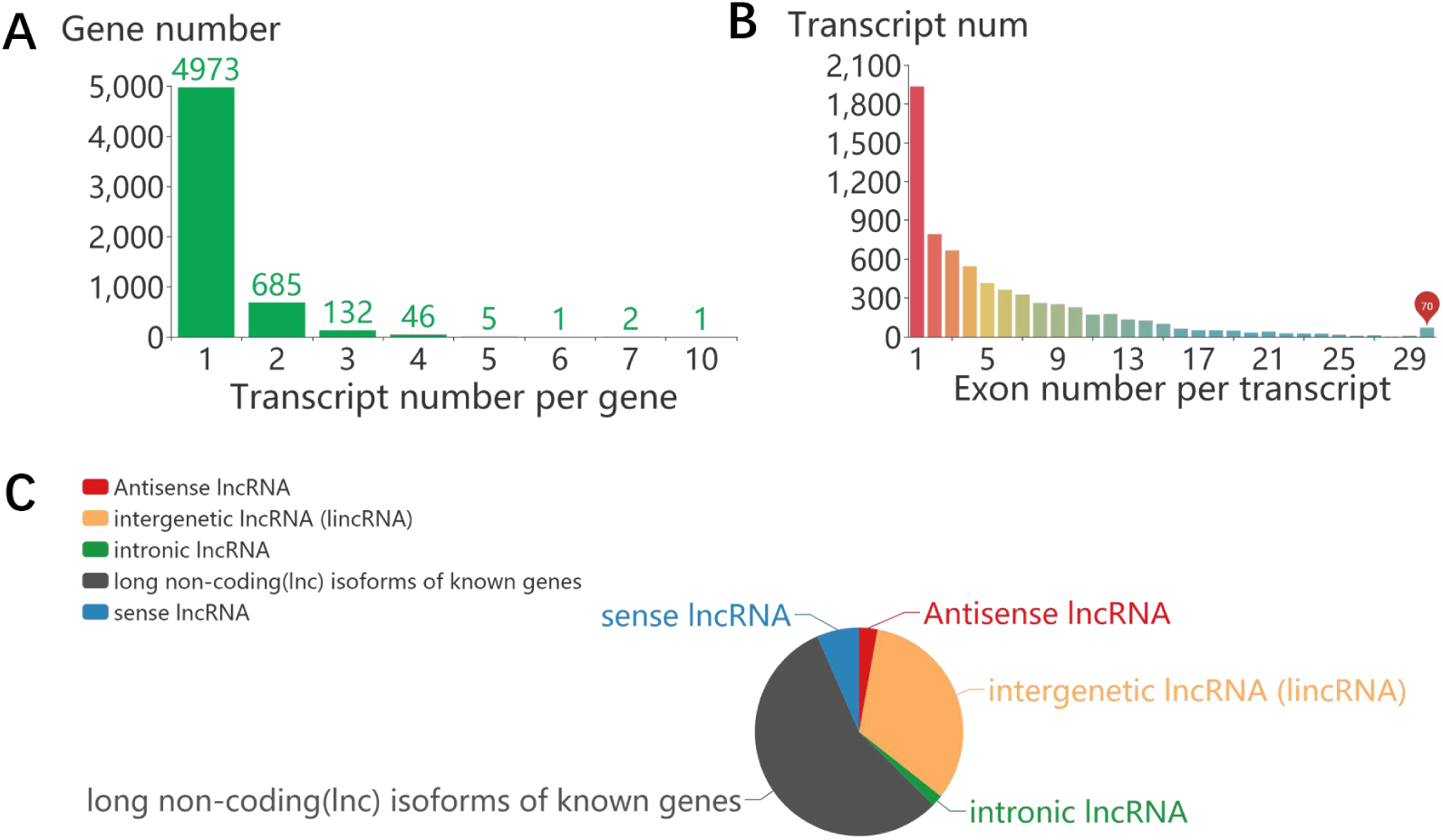
The basic molecular features of lncRNAs. A. The distribution of transcript numbers in lncRNA genes. B. The distribution of exon numbers in lncRNA transcripts. C. The classification of lncRNAs based on the positions of lncRNAs relative to protein coding genes.

### Multi-omics data integration

#### a) Expression profile

For each given lncRNA transcript, RiceLncPedia accommodates its expression profiles across all the 2313 collected RNA-seq libraries, which can be searched in expression page or downloaded in download page. Additionally, the expression of lncRNAs from a few represented projects were summarized in detail, covering diverse tissues such as leaf, stem, root, glume and panicle from Indica rice; callus, leaf, panicle before flowering, panicle after flowering, root, seed, and shoot of Japonica cv. Nipponbare, samples from various abiotic stress, such as phosphate starvation, salt stress, cadmium stress, drought stress, cold stress, osmotic stress and flood stress as well as samples grown under different hormone treatments, covering JA treatment and ABA treatment. The expression profiles can be visualized in a bar chart. Because the specific expression in a specific tissue or under a specific condition indicates the function association(38), we calculated the expression breadth, Coefficient of Variance (CV); tissue specificity index and stress responsive index (τ-Value) for each lncRNA transcript. These features greatly facilitate users to explore the lncRNAs functional associations. Users can easily search the database with CV, τ-Value, expression breadth in each selected dataset or the whole datasets (Figure 3). To explore the relationship of sample grouping with lncRNA expression profiles, we clustered the represented 339 libraries mentioned above with all 6978 lncRNAs expression values (Figure 4, Table S1). The resultant clusters were well matched between the indica and japonica groups, basically indicating the reliability of lncRNA expression profiles in RiceLncPedia. We clustered the samples into seven groups based on the hierarchical clustering heatmap. In group I, all samples belong to japonica group. The majority of tissues is root and the minor tissue is shoot. The treatments refer to JA, ABA treatment, osmotic, drought and high Cadmium stresses. Inside this group, the treatment type is a dominant feature of sub classification. Group II includes japonica leaf and shoots samples. Salt and cold treatments were grouped together, suggesting a possible similar mechanism of plant responsive salt and cold stresses. Group III covers only Japonica panicle tissues. The dominant samples in group IV are Japonica root and shoot tissues under different level of phosphate treatments. All Indica samples were classified into group V, covering different tissues such as glume, panicle, stem, leaf and root. All samples in group VI belong to Japonica shoot tissue. Although the treatments in this group refer to JA, ABA treatment, osmotic, cold, flood, drought and high Cadmium stresses and development stages, most of the developmental stages under normal conditions can still be grouped together. Some of other treatments such as JA treatment, drought, Cadmium stress samples are dispersed in developmental samples. It should be noted that these actually are the control samples in those treatments. This indicated that our lncRNA expression profiles could distinguish the control and the various treatments samples. The similar situation also occurs in group VII, where all samples belong to japonica group root tissue and the samples under control and stress conditions can be distinguished roughly. Additionally, some of different treatments such as salt, drought, cold, flood and osmotic stresses were clustered closely, indicating to some extent the similar co-expression network and plant responsive mechanisms to these stresses.

**Figure 3.**
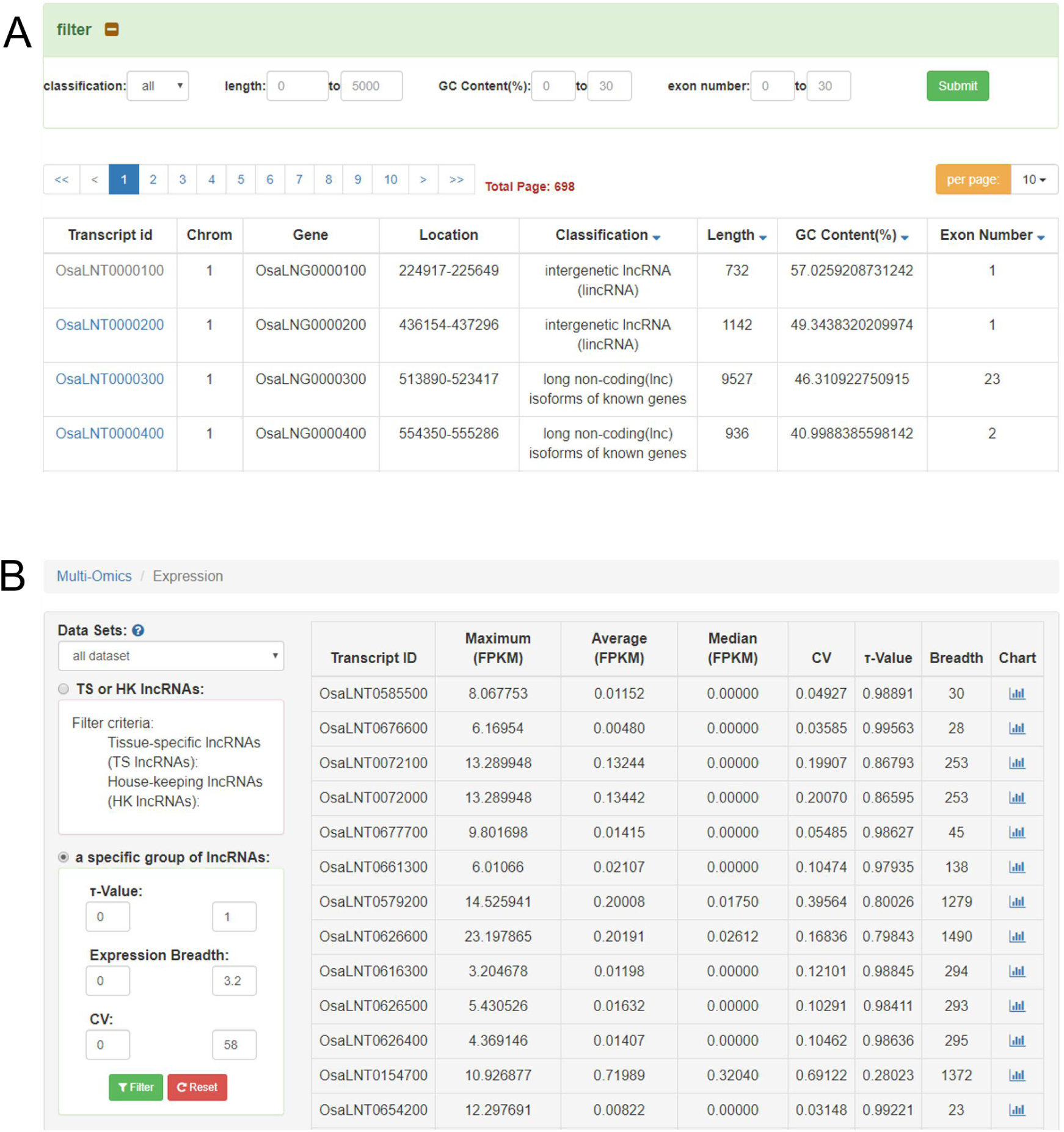
A. A snapshot of lncRNA transcripts basic molecular features. Includes transcript ids, locations, classification, length, GC content and exon numbers. B. A snapshot of lncRNA transcripts expression profile. CV, tissue Specificity Indexτ-Value and other statistics parameters were calculated for each lncRNA transcript for ten selective datasets and all datasets together.

**Figure 4.**
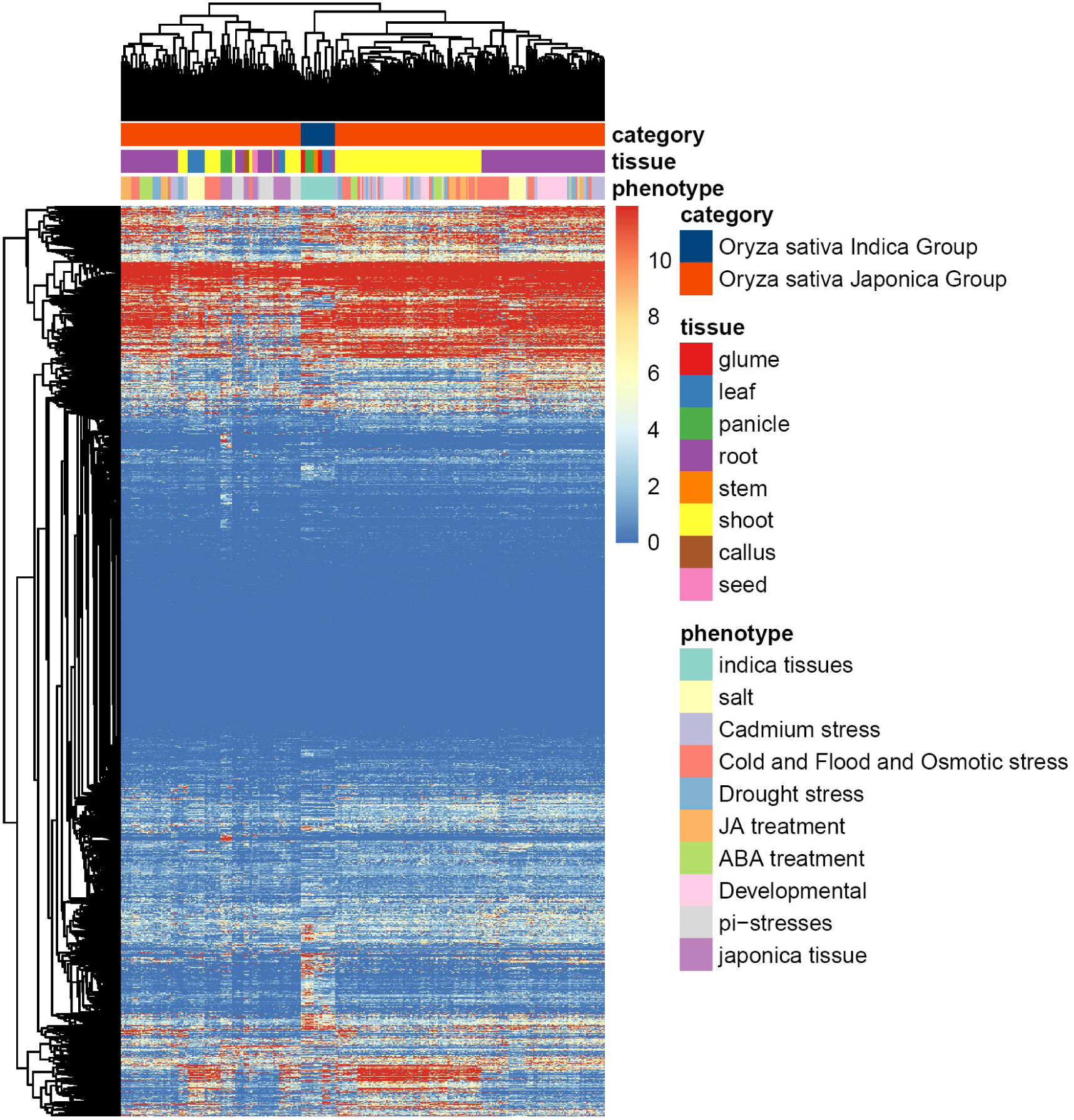
Hierarchical clustering heatmap of 339 selected RNA-seq libraries based on lncRNA expression FPKM values.In order to remove some over-expressed outliers and better distinguish between categories, we replace all expression values greater than 0.9 quantile with 0.9 quantile.

#### b) Genome variation of lncRNA

SNPs in lncRNAs were reported to be linked to lncRNA expression, structure, function and phenotypes (21). SNPs in miRNA target sites on lncRNAs may influence the miRNA-lncRNA interaction, and thereby alter their functions (44). SNPs in lncRNA transcripts have also been documented to be able to influence the lncRNA secondary structure and thereby their functions (16,45). A number of databases for SNP collection of lncRNAs in human were constructed (21,44). The databases of rice genome variation annotation linked to different features were constructed in recent years (39,46). However, the SNP annotation of lncRNAs in rice remained to be systematically collected. In RiceLncPedia, the SNPs based on 3000 genome Projects (http://snpseek.irri.org/_download.zul) were compared with the position of lncRNAs and a SNP was tagged as a lncRNA-SNP if it resides in any lncRNA. The information can promote the research of lncRNA variation association with their structures, expressions, interactions and functions. We totally identified 40758 SNPs in 5817 lncRNA transcripts with an average of about 7 SNPs per lncRNA transcript (Figure 4).

#### c) The prediction of LncRNA functions based on relative position with QTLs and GWAS tag SNPs

Trait associated SNPs were found in or nearby lncRNAs in human (47-50). In plants, an increasing number of researches have reported the regulation of agricultural traits by lncRNA-SNP identified through GWAS (16,51,52). Additionally, the co-localization of lncRNA with QTLs of complex traits has been adopted as an effective way to infer the function of lncRNAs (53-55). The fruitful information of rice GWAS and QTLs were documented recently. In RiceLncPedia, we constructed lncRNA-SNP-phenotype association if anyrice agricultural GWAS tag SNP co-located with a specific lncRNA. Similarly, a lncRNA resided in any rice QTL was also thought being association with the relevant trait. We finally found that 326 GWAS tag SNPs reside in 71 lncRNAs transcripts, which refers to 11 agricultural traits (Figure 5A). On the other hand, we found 6717 rice lncRNAs collocated with 609 traits related QTLs, such as 1000 grain weight, drought tolerance and so on, belonging to 25 tissues, development stages or stress tolerance, which are included in morphological, physiological, resistance or tolerance and other agricultural traits. The fact that most of lncRNAs (6717 out of 6978) have been co-localized in QTLs might be due to the large interval length of some QTLs. For instance, a plant height related QTL, qPH-2, identified in 2003, spans from 5263536 to 30654749 on chromosome 2 with 505 lncRNA transcripts. These QTLs were identified in earlier years, when the resolution of molecular markers, such as RFLP were not high sufficiently (Figure 5B).

**Figure 5.**
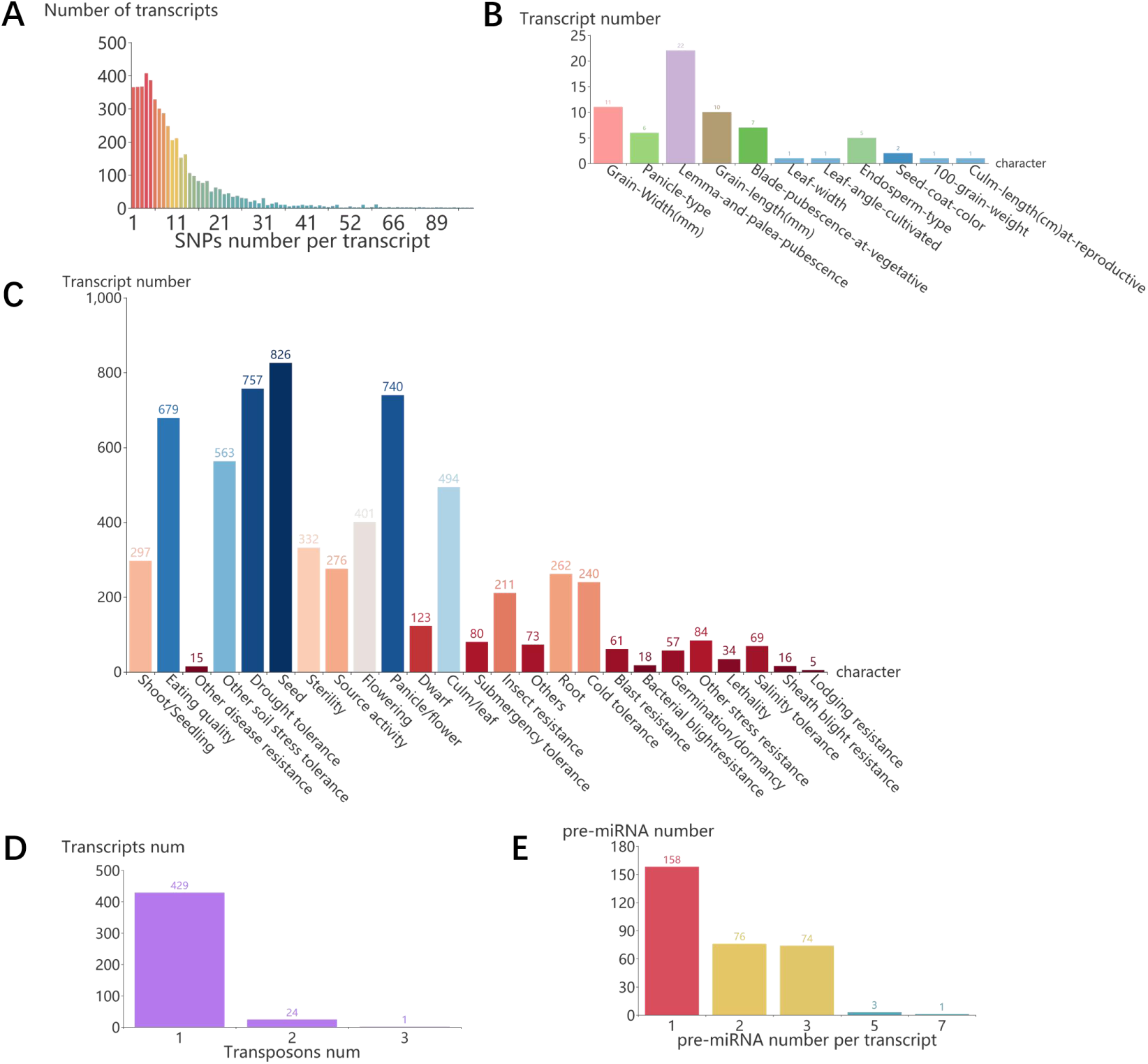
The lncRNAs features extracted from multi-omics data. A. the SNP number distribution of lncRNAs. B. The number distribution of lncRNA transcripts associated with GWAS relevant traits. C. The number distribution of lncRNA transcripts associated with QTL relevant traits. D. The number distribution of lncRNA transcripts potentially as precursors of miRNAs. E. The number distribution of lncRNA transcripts associated with transposons.

#### d) LncRNA-miRNA association

A few databases referring to the interactions of lncRNAs and miRNAs were constructed for human (21,56,57). In plants, an increasing number of studies reported lncRNAsare able to execute their roles through being targeted by specific miRNA (58-60). On the other hand, lncRNAs can act as miRNA precursors in different developmental stages in plants (61,62). To facilitate the function prediction of rice lncRNAs, the LncRNA-miRNA interactions based on the prediction by psRNA Target and those lncRNAs as precursors of miRNA were included in RiceLncPedia. In total, 6940 lncRNAs were predicted as the targets of 8184 miRNA, building up 754034lncRNA-miRNA interactions. Among of which, 6112 lncRNAs were predicted as the targets of 713 *Oraza sativa*. miRNAs, building up 65998 lncRNA and osa-miRNA interactions (Figure 6A). We also compared lncRNAs with rice miRNAs precursors and found that 335 lncRNAs have high homology with 52 pre-miRNAs, involving 590 precursor relations of lncRNAs and miRNAs (Figure 6B).

**Figure 6.**
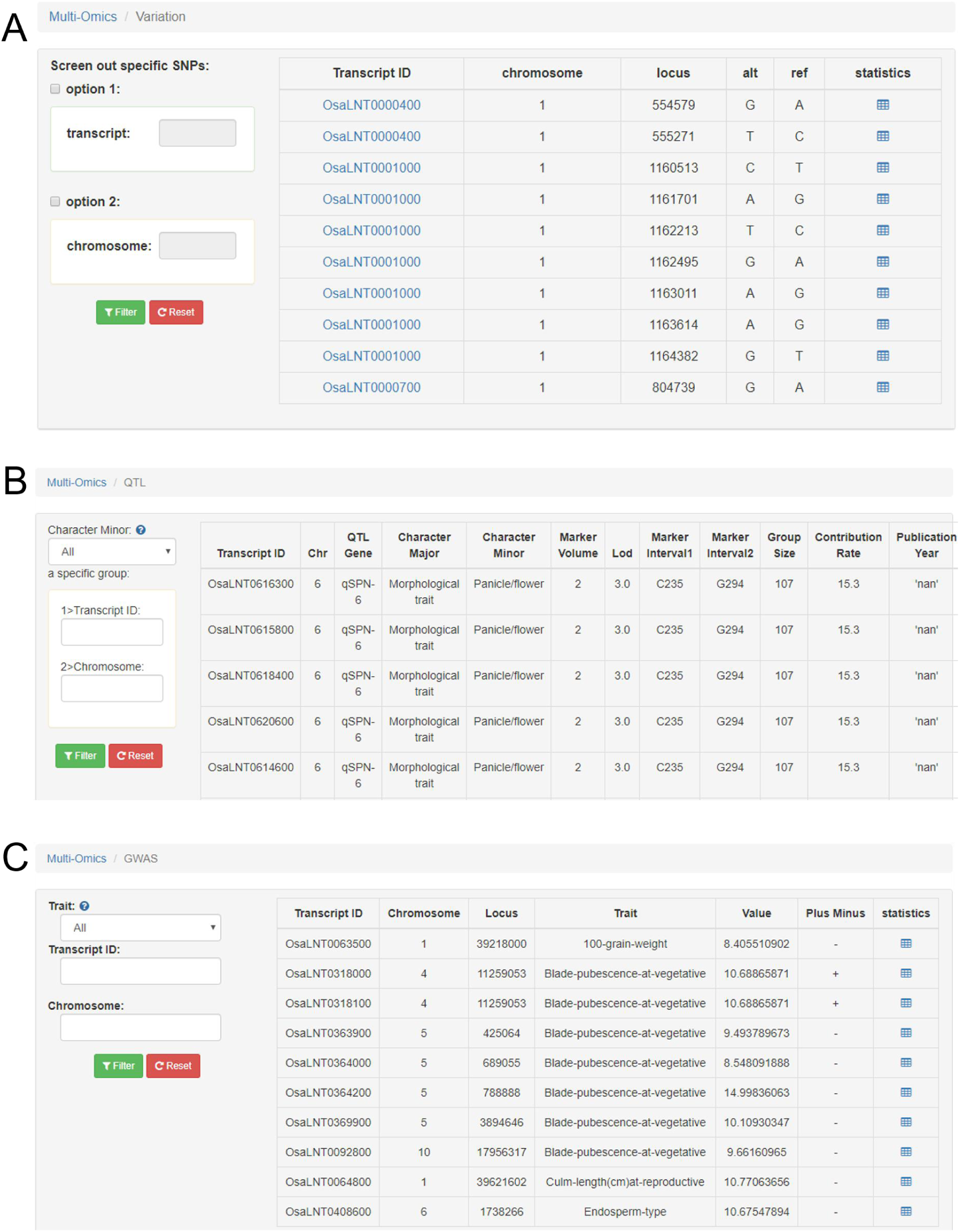
Snapshots for lncRNAs with SNP and QTL features. A. A snapshot of lncRNA transcripts encountering SNPs. B. A snapshot of lncRNA transcripts associated with agricultural traits through QTL interval positions. C. A snapshot of lncRNA transcripts associated with agricultural traits through GWAS tag SNPs.

#### e) TE-related lncRNAs

A number of lncRNAs were reported to be originated from transposons in plants and it was demonstrated that TE associated lncRNAs show tissue-specific transcription and play vital roles in plant abiotic stress responses (63,64). For this reason, LncRNAs overlapping with TE were contained in RiceLncPedia as TE-lncRNAs associations. We overall identified 380 transposons overlapped with 479 lncRNA transcripts, involving 505 transposon and lncRNA transcript relations (Figure 7).

**Figure 7.**
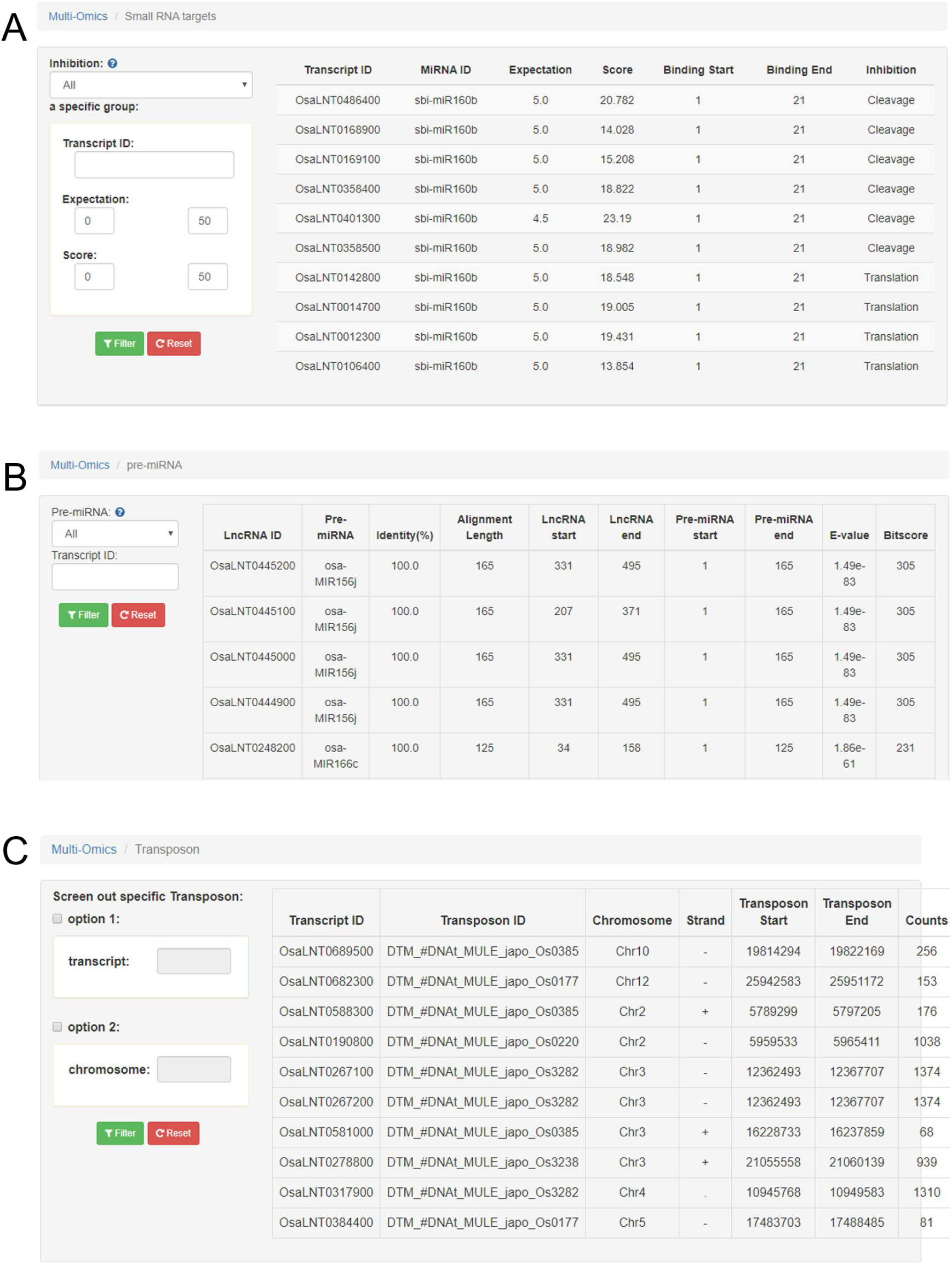
Snapshots for lncRNAs with miRNAs and TE features.A. A snapshot of lncRNA transcripts predicted as the targets. B. A snapshot of lncRNA transcripts predicted as precursors of small RNAs. C. A snapshot of lncRNA transcripts associated with transposons.

## DISCUSSION

RiceLncPedia is designed to integrate rice lncRNAs from multiple tissues under diverse conditions in a wide range of rice cultivars. Compared with the current databases containing rice lncRNAs, RiceLncPedia is more comprehensive in terms of its covered samples in which rice lncRNAs were identified and the associated lncRNA features extracted from multi-omics data, including expression profile, genome variation of lncRNA loci and association with phenotypes, LncRNA-miRNA interaction, LncRNA as potential miRNA precursors and TE-related lncRNAs. The current version of RiceLncPedia includes 6978 lncRNAs and their multifaceted genetic features. In summary, RiceLncPedia is a rich knowledge reserve of rice lncRNAs and able to serve as a valuable resource for worldwide rice research communities. Future development of RiceLncPedia will refer to regular update of newly discovered rice lncRNAs, integration of differentially expressed lncRNAs in more diverse tissues and environments, epigenetic features of lncRNAs and the association of lncRNAs with protein coding genes, experimentally validated lncRNAs and more lncRNA-phenotype associations. We are looking forward to any reasonable suggestions from worldwide scientists, with the aim to provide a continually updated and more comprehensive rice lncRNAs database.

